# IL-6 suppresses vaccine responses in neonatal mice by enhancing IL-2 activity on T follicular helper cells

**DOI:** 10.1101/2022.10.31.514554

**Authors:** Swetha Parvathaneni, Jiyeon Yang, Leda Lotspeich-Cole, Jiro Sakai, Robert C Lee, Mustafa Akkoyunlu

## Abstract

The inability of neonates to develop CD4+CXCR5^+^PD^−^1^+^ T follicular helper (T_FH_) cells contributes to their weak vaccine responses. In adult mice, IL-6 promotes T_FH_-cell expansion by suppressing the expression of IL-2Rβ on T_FH_ cells. Here, we found a totally opposite role for IL-6 in neonatal mice T_FH_ response. Whereas co-injection of neonatal mice with IL-6 and a conjugate polysaccharide vaccine suppressed T_FH_ response by increasing the production of IL-2 and expression of IL-2Rα and IL-2Rβ on T_FH_ cells, immunization of IL-6 knock-out neonatal mice led to improved antibody responses accompanied by expanded T_FH_ cells as well as lower levels of IL-2 and IL-2 receptors on T_FH_ cells. Moreover, CpG containing vaccine improved T_FH_ response in neonates while suppressing the expression of IL-2 receptors on T_FH_ cells, suggesting that CpG protects T_FH_ cells by inhibiting IL-2 activity. These findings unveil age specific differences in IL-6 mediated vaccine responses and highlight the need to consider age related immunobiological attributes in designing vaccines.

## Introduction

Newborns and infants are vulnerable to infections mostly because their immune system is not as efficient as the adult immune system in controlling microbial assaults [1; 2; 3; 4]. Early age vaccination is essential in protecting infants from infectious diseases [5; 6]. Despite the success of routine pediatric vaccination practice, neonates and infants are still vulnerable to infections because most vaccines need to be administered at 2, 4, 6 and 12 to 15 months of age in order to elicit protective immune responses [5; 6; 7; 8; 9]. Incomplete understanding of the underlying causes of suboptimal immune responses to vaccines during early age is an obstacle in improving pediatric vaccines [10]. Studies comparing neonatal and adult immune system have identified phenotypic and functional differences in the innate [11; 12] and adaptive arms [13; 14; 15; 16; 17] of the immune system between the age groups. In adult mice, generation of protective antibody responses to T cell-dependent vaccines relies on optimum germinal center reaction in secondary lymphoid organs [18]. The germinal center (GC) response involves the development of antigen specific T follicular helper (T_FH_) cells expressing the chemokine receptor CXCR5 as well as PD-1 which then migrate into B cell follicules expressing CXCL13 [18; 19; 20]. T_FH_ cells become fully committed with the expression of the transcription factor Bcl6 in response to cytokines IL-6 and IL-21. Activated T_FH_ cells interact with GC B cells which then differentiate into antibody secreting plasma cells or memory B cells [19; 21]. Tight regulation of GC reactions through promotors and inhibitors is needed to facilitate the development of high affinity antibodies against pathogens [22]. For example, FoxP3-expressing follicular regulatory T (T_FR_) cells in the GC limit T_FH_ and GC B cell responses to terminate the GC reaction [23] and also prevent autoreactive and allergen specific antibody production [24; 25; 26]. Counterbalancing the T_FH_ promoting activity of IL-21 and IL-6, cytokines IL-2 and IL-7 limit T_FH_ expansion through STAT5 activation [27; 28].

In neonates, the insufficient antibody responses to vaccines and infection are associated with inadequate expansion of T_FH_ cells and GC B cells [16; 17; 29; 30; 31]. The underlying reasons for the blunted GC response in immunized neonates are poorly understood. Our previous research indicated a predominance of T_FR_ cells in vaccinated neonatal mice resulting in higher T_FR_:T_FH_ ratio [17], which could indicate blunted GC response [26]. Importantly, we found that while IL-6 co-injection improved vaccine responses in adult mice by promoting T_FH_ expansion as shown previously [32], in neonatal mice, IL-6 co-injection suppressed T_FH_ generation and diminished antibody responses [17]. In adult mice, IL-6 protects T_FH_ cells from the well-established IL-2 mediated suppression [27] by downregulating the expression of IL-2Rβ (CD122) on T_FH_ cells and limiting IL-2 induced signaling [33].

In this study, we sought to assess how IL-6 regulated IL-2 activity on T_FH_ cells in immunized neonatal mice. We found that not only do immunized neonatal mice T_FH_ cells produced more IL-6 and IL-2 than those in adults, but they also expressed higher IL-2Rα and IL-2Rβ. Experiments performed in IL-6 co-injected wild-type neonatal and IL-6 deficient (IL-6 KO) neonatal mice indicated that, contrary to its IL-2Rβ suppressing effect in adult T_FH_ cells [33], IL-6 increased IL-2 production by T_FH_ cells and upregulated the expression of IL-2 receptors on neonatal T_FH_ cells. Moreover, immunization of neonatal mice with a CpG containing vaccine improved T_FH_ response which was accompanied by suppressed IL-6 and IL-2 production in addition to downregulated expression of IL-2 receptors on T_FH_ cells. These results provide novel insight into IL-6-mediated suppression of vaccine responses in neonatal mice.

## Results

### Enhanced IL-6 production and IL-2 activity in neonatal T_FH_ cells following immunization

We have previously shown that IL-6 co-injection suppresses vaccine responses [17]. In adult mice, IL-6 plays an important role in the improved host response to vaccines [32; 33; 34], and adjuvanted vaccines elicit peak IL-6 production in lymphoid organs during the first 24 hours after vaccination [35]. To assess the regulation of IL-6 production in neonatal splenic cells following vaccination, we immunized adult (6- to 10-week-old) and neonatal (5- to 7-day old) mice with aluminum hydroxide-adjuvanted tetanus toxoid-conjugated pneumococcal type 14 serotype (PPS14-TT) vaccine. As expected, PPS14-TT immunization induced significantly higher frequency of CD4^+^CXCR5^+^PD-1^+^FoxP3^−^ T_FH_ cells in adults than in neonates (Fig. S1A). Analysis of splenocytes by flow cytometry indicated that neonatal splenic cells contained significantly higher frequency of total (Fig. 1 A) as well as CD11c^+^ cells (Fig. 1 B) expressing IL-6 compared to adult splenocytes 24 hours after immunization. Interestingly, even substantial percentage of neonatal T cells produced IL-6 while almost no adult T cells were positive for IL-6 (Fig. S1 B). Higher neonatal IL-6 production was extended to B cells (Fig. S1 C).

**Figure 1:**
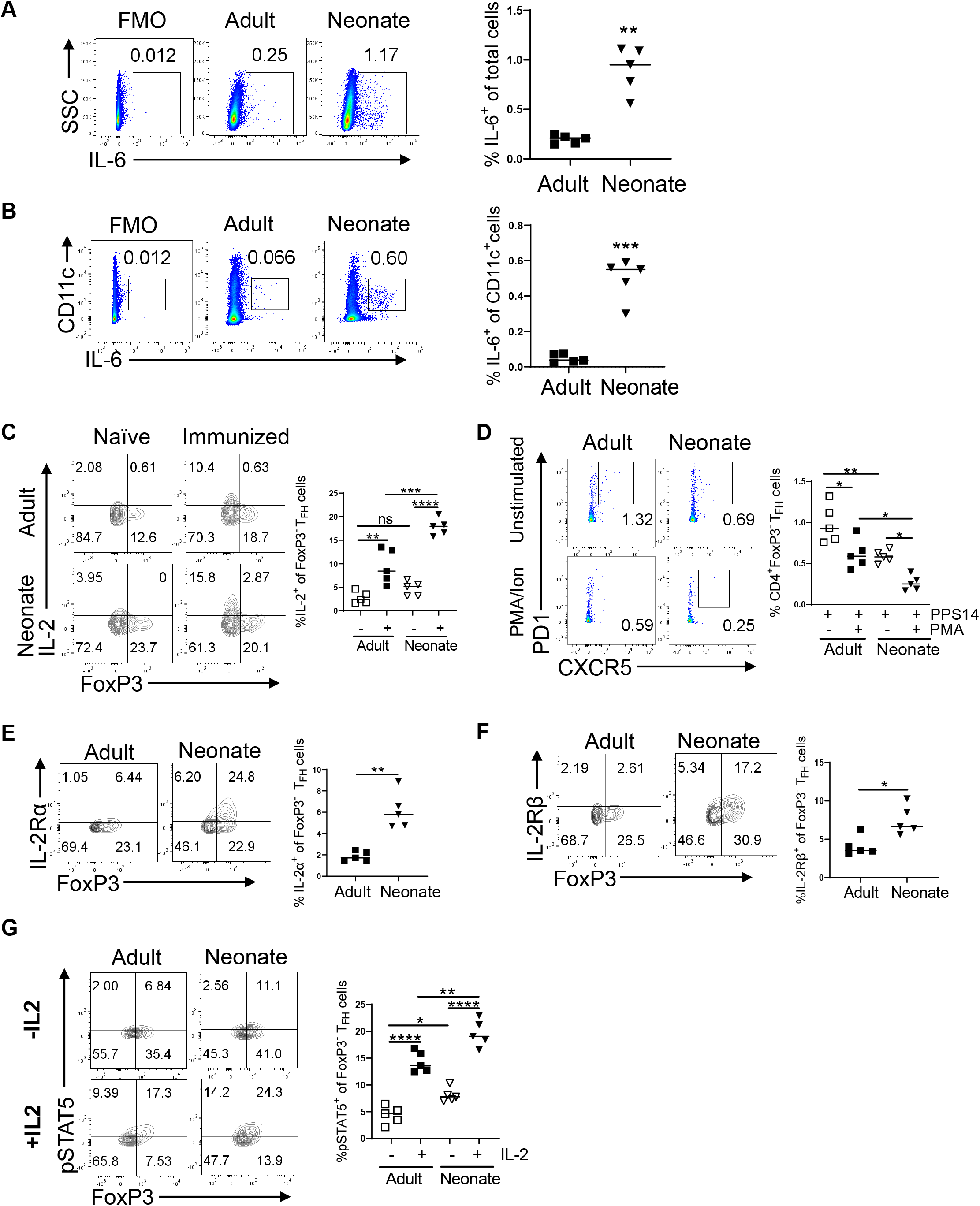
Adult (6- to 10-week-old) and neonatal (5- to 7-day-old) mice were immunized i.p. with PPS14-TT and splenocytes were analyzed by FACS. **(A)** Representative dot plots from 24 hours post immunization depict the percentages of IL-6^+^ cells pre-gated on total live cells. Mean percentages of IL-6^+^ cells among live cells are plotted (n=5). **(B)** Representative dot plots from 24 hours post immunization depict the percentages of IL-6^+^ cells pre-gated on CD11c^+^ cells. Mean percentages of IL-6^+^ cells on CD11c^+^ subsets are plotted (n=5). **(C** to **D)** Splenocytes from immunized mice were in vitro stimulated with PMA/Ion for 4 h followed by intracellular staining for IL-2 on T_FH_ cells. **(C)** Representative contour plots depict the percentages of IL-2-expressing FoxP3^+^ and FoxP3^−^ cells on T_FH_ cells. Mean percentages of IL-2^+^ cells among FoxP3^−^ T_FH_ cells are plotted (n=5). **(D)** Representative dot plots depict the percentage of T_FH_ (CXCR5^+^PD1^+^) cells pre-gated on CD4^+^Foxp3^−^ cells. Mean percentages of T_FH_ cells are plotted (n=5). Representative contour plots depict percentages of IL-2Rα-**(E)** or IL-2Rβ-**(F)** expressing FoxP3^+^ and FoxP3^−^ cells on T_FH_ cells. Mean percentages of IL-2Rα^+^ and IL-2Rβ^+^ cells among FoxP3^−^ T_FH_ cells are also plotted (n=5). **(G)** Splenocytes from 7 dpi were stimulated with or without recombinant IL-2 for 15 minutes, followed by intracellular staining for p-STAT5. Representative contour plots depict the percentages of p-STAT5^+^ cells among FoxP3^+^ and FoxP3^−^ T_FH_ (pre-gated on CD4^+^CXCR5^+^PD1^+^) cells. Mean percentages of p-STAT5^+^ cells among FoxP3^−^ T_FH_ cells are plotted (n=5). The data are representative of at least two independent experiments. Each experiment was performed twice.

Next, we focused on IL-2 production in immunized mice because a recent report demonstrated that FoxP3^−^CD4^+^ T cells spontaneously produced significantly more IL-2 in unimmunized neonatal mice compared to adults and suggested that the elevated IL-2 production likely contributes to the ablated T_FH_ development in immunized neonatal mice [31]. Consistent with this report, we found higher IL-2 expression by freshly isolated naïve neonatal CD4^+^ T cells than the naïve adult counterparts (Fig. S1 D). The proportion of neonatal CD4^+^IL-2^+^ cells compared to those of adults increased further seven days after immunization as assessed by ex vivo staining (Fig. S1 D) or following phorbol myristate acetate/ionomycin (PMA/Ion) stimulation (Fig. S1 E). We also measured the frequencies of IL-2^+^ T_FH_ cells in adult and neonatal mice because Papillon and colleagues showed that the expression of IL-2 by T_FH_ cells is more relevant for T_FH_ suppression than IL-2 produced by other cells [33]. As with total CD4^+^ cells, naïve neonatal T_FH_ cells expressed more IL-2 than their adult counterparts and this difference increased further after immunization when measured ex vivo (Fig. 1 C) or following PMA/Ion stimulation (Fig. S1F). The increase in IL-2 production by T_FH_ cells following PMA/Ion stimulation of cells was meaningful because there was a significant decrease in the frequency of neonatal and adult T_FH_ cells in parallel to the increase in IL-2 production (Fig. 1 D).

In addition to the amount of IL-2 produced by T_FH_ cells, the permissiveness T_FH_ cells to IL-2 mediated suppression also depends on the level of IL-2 receptor expression by T_FH_ cells [33], and in adult mice IL-6 alleviates IL-2 mediated T_FH_ suppression by downregulating IL-2Rβ expression on T_FH_ cells [33]. We therefore compared IL-2Rα and IL-2Rβ expression on adult and neonatal T_FH_ cells following immunization. We found significantly higher expression of IL-2Rα and IL-2Rβ on neonatal T_FH_ cells compared to adult cells (Fig. 1 E and F). Thus, not only did neonatal T_FH_ cells produce more IL-6 and IL-2 than the adult cells, but neonatal T_FH_ cells express higher levels of both IL-2 receptors than adult T_FH_ cells. The T_FH_ inhibitory activity of IL-2 is mediated by STAT5 [36]. To test whether elevated IL-2Rα and IL-2Rβ resulted in enhanced STAT5 activation in T_FH_ cells, we isolated splenocytes seven days post immunization (dpi) and subjected them to IL-2 stimulation. We found that even unstimulated neonatal FoxP3^−^ T_FH_ cells contained significantly higher phospho-STAT5^+^ (p-STAT5^+^) cells than those in adult cells (Fig. 1 G). More importantly, correlating with the increase in IL-2Rα and IL-2Rβ expression, IL-2 stimulation induced significantly higher p-STAT5+ T_FH_-cell-frequency in immunized neonates than in adults (Fig. 1 G). Since IL-2 induced STAT5 activation blunts the expression of *Bcl6* [27; 36; 37], diminished T_FH_ development in neonatal mice is likely due to elevated IL-2Rα and IL-2Rβ expression on T_FH_ cells in addition to increased IL-2 production by T_FH_ cells.

### Co-injection of IL-6 suppresses vaccine responses in neonates by enhancing IL-2 activity

In adult mice, IL-6 enhances vaccine responses by promoting T_FH_ cells [17; 32] through downregulation of IL-2Rβ expression and protecting T_FH_ cells from the suppressive effect of IL-2 [33]. We previously found that IL-6 co-injection with PPS14-TT vaccine suppresses T_FH_ development and antibody responses in neonatal mice [17]. To assess whether IL-6 mediated blunting of PPS14-TT response involves the regulation of IL-2 activity, we co-injected adult and neonatal mice with IL-6 and PPS14-TT vaccine or PPS14-TT vaccine in PBS and analyzed the changes in IL-2 production and the expression of IL-2 receptors on T_FH_ cells. Confirming our previous results [17], IL-6 co-injection decreased T_FH_ cell frequency in neonatal mice, whereas it increased T_FH_ population in adult mice (Fig. 2 A). Also as we showed before, there was a significant increase in FoxP3+CD4+PD-1+CXCR5+ T_FR_ cells in IL-6 co-injected neonatal mice while adult T_FR_ cells decreased with excess IL-6 (Fig. S2 A). Interestingly, the decrease in T_FH_ cells was accompanied by a reciprocal increase in IL-2 expressing CD4+ as well as FoxP3^−^ T_FH_ cells in IL-6 co-injected neonatal mice compared to neonates that received PPS14-TT alone (Fig. S2 B and C). Conversely, IL-6 co-injection with PPS14-TT led to a significant decrease in IL-2 producing CD4+ and FoxP3^−^ T_FH_ cells in adults compared to those injected with PPS14-TT alone. In vitro stimulation of splenocytes from immunized neonatal mice with PMA/Ion further increased the IL-2 production from CD4+ and T_FH_ cells (Fig. 2S D and Fig. 2 B). In parallel to the increase in IL-2^+^ T_FH_ cell population following PMA/Ion stimulation, there was a statistically significant decrease in the FoxP3^−^ T_FH_ population in IL-6 co-injected neonates compared to those given the vaccine only (Fig. 2 C). Moreover, consistent with the previous report [33], we found a decrease in the frequency of IL-2Rβ-expressing (Fig. 2 E), but not IL-2Rα-expressing (Fig. 2 D), T_FH_ cells in IL-6 co-injected adult mice compared to those injected with PPS14-TT alone. The decrease in IL-2Rβ expression was biologically relevant because stimulation of splenocytes from immunized adult mice led to a decrease in the frequency of p-STAT5^+^ T_FH_ cells from IL-6 co-injected mice compared to those immunized with PPS14-TT alone (Fig. 2 F). In sharp contrast to adult mice, IL-6 co-injection increased both IL-2Rα and IL-2Rβ expression on neonatal T_FH_ cells (Fig. 2 D and E). Accompanying the increase in both the IL-2 receptors, IL-6 co-injected neonatal mice T_FH_ cells manifested higher frequency of p-STAT5^+^ population as compared to mice injected with PPS14-TT alone following IL-2 stimulation of splenocytes harvested seven dpi (Fig. 2 F). Thus, the decreased IL-2 production by T_FH_ cells together with the dampened IL-2Rβ expression on T_FH_ cells are likely responsible for the increased T_FH_ cell frequency in IL-6 co-injected adult mice. Paradoxically, in neonates, excess IL-6 further increases the production of IL-2 by T_FH_ cells and stimulates the expression of both the IL-2 receptors on T_FH_ cells, likely rendering them more susceptible to IL-2 mediated suppression.

**Figure 2:**
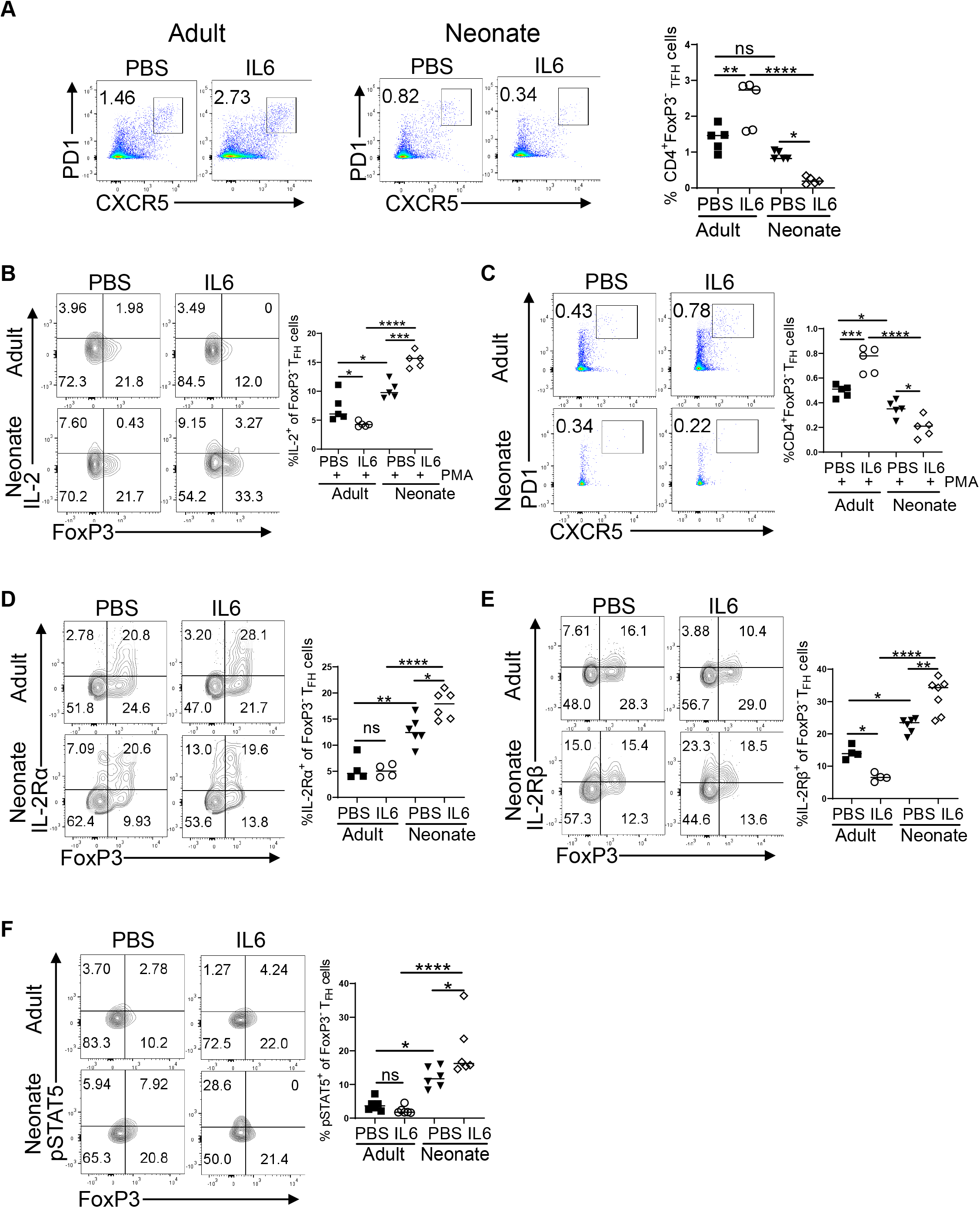
Adult (6- to 10-week-old) and neonatal (5- to 7-day-old) mice were immunized i.p. with PPS14-TT in PBS (PBS) or PPS-14-TT + IL-6 (IL6) and splenocytes were analyzed by flow cytometry at 7 dpi. **(A)** Representative dot plots depict the percentages of T_FH_ (CXCR5^+^PD1^+^) cells pre-gated on CD4^+^FoxP3^−^ cells. Mean percentages of T_FH_ cells are plotted (n=5). **(B and C)** Splenocytes from immunized mice were in vitro stimulated with PMA/Ion for 4 h followed by intracellular staining for IL-2 on T_FH_ cells. **(B)** Representative contour plots depict the percentages of IL-2-expressing FoxP3^+^ and FoxP3^−^ cells on T_FH_ cells. Mean percentages of IL-2^+^ cells among FoxP3^−^ T_FH_ cells are plotted (n=5). **(C)** Representative dot plots depict the percentage of T_FH_ (CXCR5^+^PD1^+^) cells pre-gated on CD4^+^Foxp3^−^ cells. Mean percentages of T_FH_ cells are plotted (n=5). **(D and E)** Splenocytes from immunized mice were pre-gated on CD4^+^CXCR5^+^PD1^+^ T_FH_ cells. Representative contour plots depict percentages of IL-2Rα- **(D)** or IL-2Rβ- **(E)** expressing FoxP3^+^ and FoxP3^−^ cells on T_FH_ cells. Mean percentages of IL-2Rα^+^ and IL-2Rβ^+^ cells among FoxP3^−^ T_FH_ cells are also plotted (n=4-7). **(F)** Splenocytes from immunized mice were stimulated with recombinant IL-2 for 15 minutes, followed by intracellular staining for pSTAT5. Representative contour plots depict the percentage of pSTAT5^+^ cells on T_FH_ (CXCR5^+^PD1^+^Foxp3^−^) cells pre-gated on CD4^+^ cells. Mean percentages of pSTAT5^+^ cells among T_FH_ cells are plotted (n=6). Experiments were performed two to four times.

### IL-6 deficiency improves vaccine responses in neonatal mice

Immunization of neonatal C57BL/6 mice indicated that elevated IL-6 together with increased IL-2 production and upregulated IL-2Rα and IL-2Rβ may be preventing the expansion of T_FH_ cells (Fig. 1). IL-6 co-injection studies suggested a role for IL-6 in the induction of IL-2 and in the increase of IL-2Rα and IL-2Rβ expression by T_FH_ cells, accompanied by elevated p-STAT5^+^ T_FH_ cells (Fig. 2). Collectively, these data support a role for IL-6 in IL-2 mediated suppression of T_FH_ generation in neonates (Fig. 2 A) [17]. To test this hypothesis, we immunized wild-type and IL-6 KO neonatal mice with PPS14-TT vaccine and characterized the immune responses. We found that IL-6 KO mice mounted significantly more anti-PPS14 IgG antibodies at 30 days post-immunization compared to wild-type neonates (Fig. 3 A). In parallel to the increase in antibody responses, the frequencies of both the B220^+^GL7^+^Fas^+^ GC B cell and the CD4^+^PD-1^+^CXCR5^+^FoxP3^−^ T_FH_ cell populations were significantly higher in IL-6 KO mice than those in wild-type mice (Fig. 3 B and C). Our IL-6 co-injection study had resulted in an increase in the CD4^+^PD-1^+^CXCR5^+^FoxP3^+^ T_FR_ population reciprocal to the decrease in T_FH_ cell frequency (Fig. S2 A) [17]. In support of a role for IL-6 in promoting T_FR_ cells, we measured significantly lower frequency of T_FR_ cells in immunized IL-6 KO mice compared to wild-type mice (Fig. 3 D).

**Figure 3:**
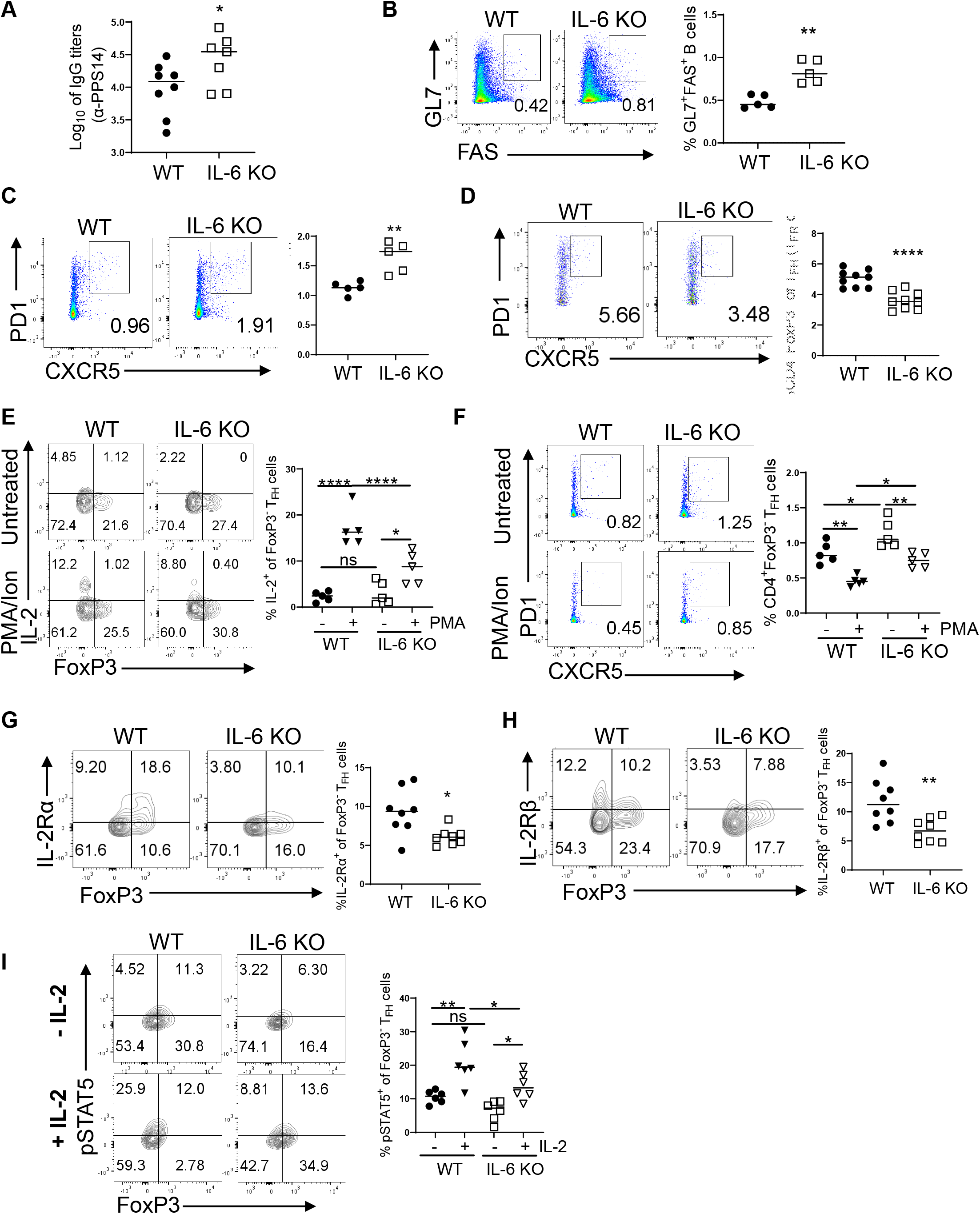
Neonatal (5- to 7-day-old) wild-type (C57BL/6J) and IL-6 KO mice were immunized i.p. with PPS14-TT and splenocytes were analyzed by flow cytometry at 7 dpi. **(A)** Serum anti-PPS14 IgG titers were determined by ELISA 4 weeks after immunization (n=5). **(B)** Representative dot plots depict the GC (GL7^+^FAS^+^) cells pre-gated on B220^+^ cells. Mean percentages of GC B cells are plotted (n=5). **(C)** Representative dot plots depict the GC T_FH_ (CXCR5^+^PD1^+^) cells pre-gated on CD4^+^FoxP3-cells. Mean percentages of FoxP3^−^ T_FH_ cells are plotted (n=5). **(D)** Representative dot plots depict the GC T_FR_ (CXCR5^+^PD1^+^) cells pre-gated on CD4^+^FoxP3^+^ cells. Mean percentages of FoxP3^+^ T_FR_ cells are plotted (n=9). **(E and F)** Splenocytes from immunized mice were in vitro stimulated with PMA/Ion for 4 h and T_FH_ cells were analyzed. **(E)** Representative contour plots depict the percentages of IL-2-expressing FoxP3^+^ and FoxP3-cells pre-gated on T_FH_ (CD4^+^CXCR5^+^PD1^+^) population. Mean percentages of IL-2^+^ cells among Foxp3^−^ T_FH_ cells are plotted (n=5). **(F)** Representative dot plots depict the percentages of T_FH_ (CXCR5^+^PD1^+^) cells pre-gated on CD4^+^Foxp3^−^ cells. Mean percentages of Foxp3^−^ T_FH_ cells are plotted (n=5). **(G and H)** Splenocytes from immunized mice were pre-gated on CD4^+^CXCR5^+^PD1^+^ T_FH_ cells. Representative contour plots depict percentages of IL-2Rα- **(G)** or IL-2Rβ- **(H)** expressing FoxP3^+^ and FoxP3-cells on T_FH_ cells. Mean percentages of IL-2Rα^+^ and IL-2Rβ^+^ cells among FoxP3^−^ T_FH_ cells are also plotted (n=4-5). **(I)** Splenocytes from immunized mice were stimulated with or without recombinant IL-2 for 15 minutes, followed by intracellular staining for pSTAT5. Representative contour plots depict the percentages of pSTAT5^+^ cells among FoxP3^+^ and FoxP3^−^ T_FH_ (pre-gated on CD4^+^CXCR5^+^PD1^+^) cells. Mean percentages of pSTAT5^+^ cells among FoxP3^−^ T_FH_ cells are plotted (n=6). Experiments were performed two to seven times.

Next, we focused on IL-2 activity in the absence of IL-6 in immunized neonatal mice. We began by comparing the production of IL-2 in immunized IL-6 KO and wild-type neonates. At seven dpi there was no difference in the frequency of CD4^+^IL-2^+^ cell population between the two mouse strains (Fig. S2 E). However, in IL-6 KO mice CD4^+^PD-1^+^CXCR5^+^FoxP3^−^ T_FH_ cell population contained significantly less IL-2^+^ T_FH_ cells than the wild-type mice (Fig. S2 F). Like the ex vivo analyzed cells, in vitro stimulation of splenocytes from immunized neonatal mice with PMA/Ion also showed no difference in IL-2-producing CD4^+^ population between wild-type and IL-6 KO mice (Fig. S2 G) but there was a significant decrease in IL-2^+^ T_FH_ cell frequency in IL-6 KO mice (Fig. 3 E). Importantly, in parallel to the decrease in IL-2^+^ T_FH_ population, PMA/Ion stimulation resulted in higher frequency of T_FH_ population in IL-6 KO mice than the wild-type mice (Fig. 3 F).

The absence of systemic IL-6 also led to a change in the expression of IL-2 receptors on T_FH_ cells: both the receptors were downregulated on T_FH_ cells from immunized IL-6 KO mice compared to wild-type mice (Fig. 3 G and H). The decreases in the expression of IL-2 receptors on T_FH_ cells were biologically meaningful because in vitro stimulation of splenocytes from immunized mice with IL-2 resulted in a significantly reduced frequency of p-STAT5^+^ T_FH_ cells in IL-6 KO mice than those from wild-type mice (Fig. 3 I). Taken together, unlike in adult mice [33], neonatal mice responses to vaccines improve in the absence of IL-6. Both the ablation of IL-2 mediated suppression and the blunting of T_FR_ response likely contribute to the improved vaccine responses in neonatal IL-6 KO mice.

### CpG enhances vaccine response in neonatal mice through suppression of IL-6 and IL-2 signaling in T_FH_ cells

The IL-6 co-injection and IL-6 KO mice studies highlighted the critical role for IL-6 in suppressing neonatal T_FH_ generation through the modulation of IL-2 activity. CpG improves vaccine responses in neonates by stimulating T_FH_ and GC B cell responses [16]. To test whether the improvement by CpG also involves the modulation of IL-6 and IL-2 activity, we characterized the immune responses in neonates immunized with CpG containing PPS14-TT or PPS14-TT vaccine in PBS. As shown previously [16], CpG significantly increased serum IgG1 and IgG2c antibody levels against PPS14 (Fig. 4 A). CpG also increased the frequencies of GC B (Fig. 4 B) and T_FH_ cells (Fig. 4 C) in spleens of immunized neonates (Fig. 4B and C). Moreover, the increased frequency of T_FH_ cells was accompanied by decreased frequency of T_FR_ cells (Fig. 4 D) and T_FR_:T_FH_ ratio (Fig. 4 E).

**Figure 4:**
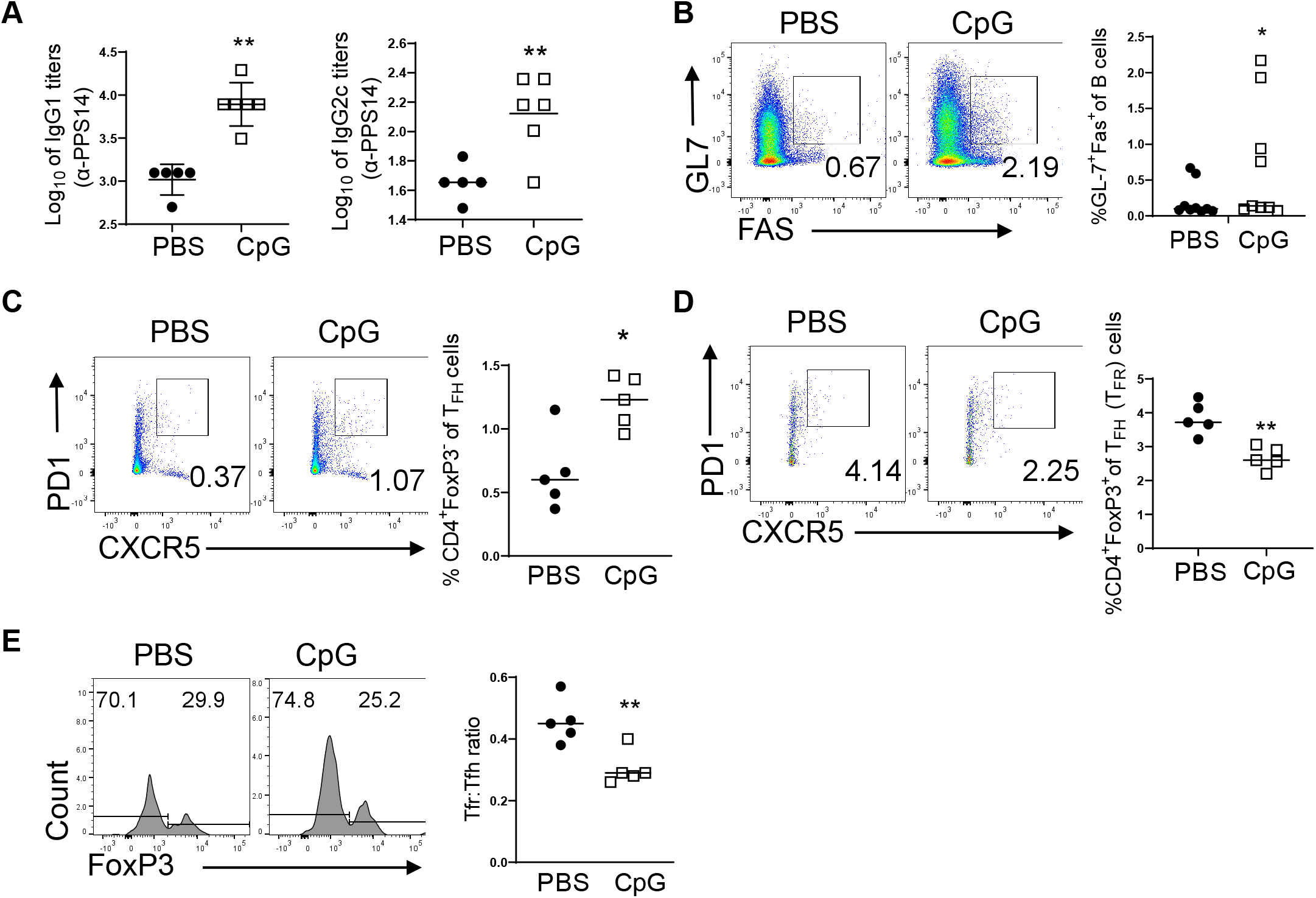
C57BL/6J (5- to 7-day-old) mice were immunized i.p. with PPS14-TT (PBS) or PPS14-TT + CpG (CpG) and splenocytes were analyzed by flow cytometry at 7 dpi. **(A)** Serum anti-PPS14 IgG1 and anti-PPS14 IgG2c titers were determined by ELISA four weeks after immunization (n=5). **(B)** Representative dot plots depict the GC (GL7^+^FAS^+^) cells pre-gated on B220^+^ cells. Mean percentages of GC B cells are plotted (n=9). **(C)** Representative dot plots depict the GC T_FH_ (CXCR5^+^PD1^+^) cells pre-gated on CD4^+^FoxP3^−^ cells. Mean percentages of T_FH_ cells are plotted (n=5). **(D)** Representative dot plots depict the GC T_FR_ (CXCR5^+^PD1^+^) cells pre-gated on CD4^+^FoxP3^+^ cells. Mean percentages of T_FR_ cells are plotted (n=5). **(E)** Histograms represent the T_FH_ (FoxP3^−^) and T_FR_ (FoxP3^+^) populations on CD4+CXCR5^+^PD1^+^ gated cells and the ratio of T_FR_ to T_FH_ cells (T_FR_:T_FH_) are plotted (n=5). Experiments were performed two to eight times.

Next, we measured the production of IL-6 following immunization. The percentages of IL-6^+^ splenocytes as well as CD11c^+^ cells were significantly lower in neonates immunized with CpG containing vaccine 24 hours after vaccination (Fig. 5 A and B). Ex vivo staining of freshly isolated cells seven dpi indicated that CpG containing PPS14-TT vaccine elicited significantly lower frequencies of IL-2-expressing CD4^+^ and CD4^+^PD-1^+^CXCR5^+^FoxP3^−^ T_FH_ cells than those immunized with PPS14-TT alone (Fig. S3 A and SB). As with ex vivo stained cells, PMA/Ion stimulation increased the CD4^+^IL-2^+^ population as well as the IL-2^+^ T_FH_ cells compared to unstimulated cells from both the immunized groups. However, the frequencies of IL-2^+^ cells from both the CD4^+^ (Fig. S3 C) and T_FH_ cells (Fig. 5 C) were significantly lower in neonates immunized with CpG containing PPS14-TT vaccine compared to those immunized with PPS14-TT alone. More importantly, the increase in IL-2 production directly correlated with a reciprocal decrease in the frequency of T_FH_ cells (Fig. 5 D). Lower production of IL-2 in neonates immunized with the CpG containing vaccine translated into higher frequency of T_FH_ cells than neonates immunized with PPS14-TT alone.

**Figure 5:**
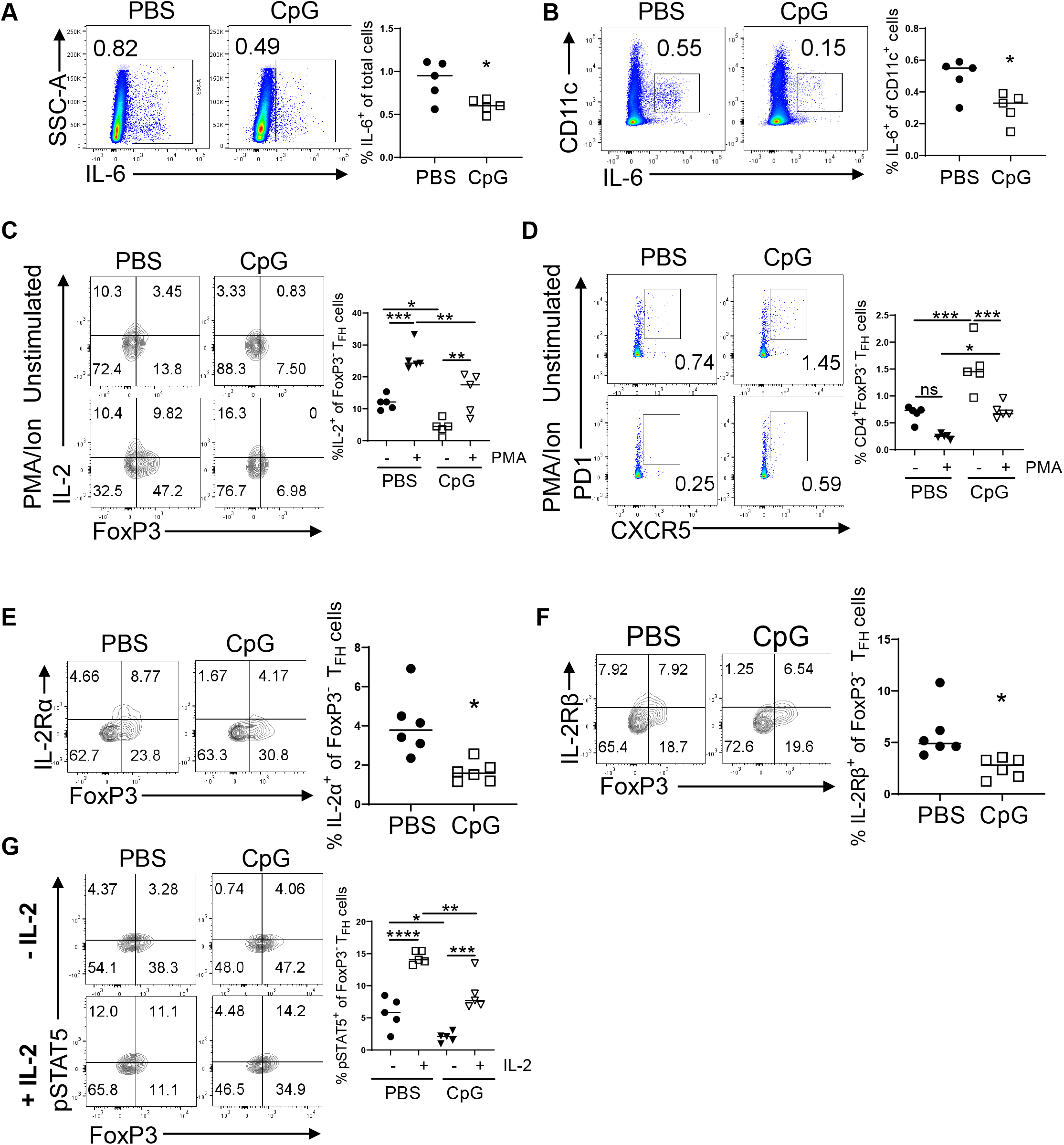
C57BL/6J (5- to 7-day-old) mice were immunized i.p. with PPS14-TT (PBS) or PPS14-TT + CpG (CpG) and splenocytes were analyzed by flow cytometry. **(A)** Representative dot plots from 24 hours post immunization depict the percentages of IL-6^+^ cells pre-gated on total live cells. Mean percentages of IL-6^+^ cells among live cells are plotted (n=5). **(B)** Representative dot plots from 24 hours post immunization depict the percentages of IL-6^+^ cells pre-gated on CD11c^+^ cells. Mean percentages of IL-6^+^ cells on CD11c^+^ subsets are plotted (n=5). **(C and D).** Splenocytes from 7 dpi were in vitro stimulated with PMA/Ion for 4 h and T_FH_ cells were analyzed. **(C)** Representative contour plots depict the percentages of IL-2^+^ cells among T_FH_ (CXCR5^+^PD1^+^) population pre-gated on CD4^+^Foxp3^−^ cells. Mean percentages of IL-2^+^ cells among Foxp3^−^ T_FH_ cells are plotted (n=5). **(D)** Representative dot plots depict the percentages of T_FH_ (CXCR5^+^PD1^+^) cells pre-gated on CD4^+^Foxp3^−^ cells. Mean percentages of Foxp3^−^ T_FH_ cells are plotted (n=5). **(E and F)** Splenocytes from 7 dpi were pre-gated on CD4^+^CXCR5^+^PD1^+^ T_FH_ cells. Representative contour plots depict percentages of IL-2Rα **(E)** or IL-2Rβ **(F)** expressing FoxP3^+^ and FoxP3^−^ cells on T_FH_ cells. Mean percentages of IL-2Rα^+^ and IL-2Rβ^+^ cells among FoxP3^−^ T_FH_ cells are also plotted (n=5). **(G)** Splenocytes from 7 dpi were stimulated with or without recombinant IL-2 for 15 minutes, followed by intracellular staining for pSTAT5. Representative contour plots depict the percentages of pSTAT5^+^ cells among FoxP3^+^ and FoxP3^−^ T_FH_ (pre-gated on CD4^+^CXCR5^+^PD1^+^) cells. Mean percentages of pSTAT5^+^ cells among FoxP3^−^ T_FH_ cells are plotted (n=5). Experiments were performed twice.

Inclusion of CpG in PPS14-TT vaccine also influenced the expression of IL-2 receptors. Both, IL-2Rα^+^ and IL-2Rβ^+^ T_FH_ cell frequencies were significantly lowered by CpG (Fig. 5 E and F). The decrease in the expression of IL-2 receptors had functional consequence because stimulation of splenocytes from CpG containing vaccine led to lower frequencies of p-STAT5^+^ T_FH_ cells than those immunized with PPS14-TT alone following the stimulation of purified splenocytes with IL-2 (Fig. 5 G). Thus, CpG improves vaccine responses by protecting T_FH_ cells from the suppressive activity of IL-2 through the reduction of IL-6 and IL-2 production and by decreasing IL-2Rα and IL-2Rβ expression on T_FH_ cells.

## Discussion

The inability of neonates to develop fully mature GC reaction is mostly responsible for their suboptimal vaccine responses [7; 16; 17; 30]. We previously reported suppression of neonatal T_FH_ and antibody responses when IL-6 is co-injected with PPS14-TT vaccine [17]. This is in sharp contrast to adult mice, which mount increased antibody [17; 32] and T_FH_ [17] responses in the presence of access IL-6. Here, we showed that excess IL-6 sensitizes T_FH_ cells to the suppressive cytokine IL-2 by increasing the expression of IL-2 receptors on T_FH_ cells and by inducing the production of IL-2 by T_FH_ cells. Conversely, vaccine response is improved in neonatal IL-6 KO mice. The improvement in IL-6 KO mice is accompanied by dampened IL-2 production and reduced IL-2 receptor expression on T_FH_ cells. Underscoring this unique mechanism, we showed that inclusion of CpG in vaccine improved T_FH_ response in addition to decreasing IL-6 and IL-2 production and suppressing IL-2Rα and IL-2Rβ expression on T_FH_ cells.

The suppressive activity of IL-2 on T_FH_ cells is well established [18; 27]. In the current study, we focused on the regulation of IL-2 mediated suppression of neonatal T_FH_ cells because Papillion and colleagues reported that in adult mice the expansion of T_FH_ cells is dependent on IL-6 mediated downregulation of IL-2Rβ on T_FH_ cells and limiting STAT5 phosphorylation [33]. We first observed that immunized neonatal mice spleens contained significantly more IL-6 producing CD11c^+^, CD19^+^ and CD3^+^ cells than the adult mice. In a recent study, Pyle and colleagues reported increased IL-2 production in neonatal pulmonary lymph nodes following respiratory syncytial virus infection [31]. Highlighting the role for IL-2 in the suppression of T_FH_ cells, they found that inhibition of IL-2 led to expanded T_FH_ population in infected neonatal mice, as was shown previously in adult mice [36; 38]. These authors also showed higher expression of IL-2Rα and IL-2Rβ on naïve CD44^−^CD62L^+^FoxP3^−^CD4^+^ cells and elevated p-STAT5 activity in CD62L^+^FoxP3^−^ CD4^+^ cells following in vitro stimulation with IL-2. However, they did not assess the regulation of IL-2 receptor expression and p-STAT5 activation in infected or immunized neonatal mice T_FH_ cells and they concluded that the increased receptor expression in naïve neonatal mice CD4^+^ cells is unlikely to play a role in heightened IL-2 activity in infected neonatal mice CD4^+^ cells from pulmonary lymph nodes. Like reported by Pyle and colleagues [31], we also measured higher expression of IL-2 by naïve and immunized neonatal mice CD4^+^ cells. In a recent study, Papillion and colleagues showed that IL-2 produced by T_FH_ cells is especially instrumental for the suppression of T_FH_ cells [33]. Further gating of CD4^+^ cells for FoxP3^−^PD-1^+^CXCR5^+^ T_FH_ cells indicated that IL-2 production was especially elevated in this population following the immunization of neonates. Moreover, we measured higher frequencies of IL-2Rα and IL-2Rβ expressing T_FH_ cells in neonatal mice compared to adult mice following immunization. The increase in IL-2 receptor positive T_FH_ cells in immunized neonatal mice was biologically relevant because in vitro stimulation of CD4^+^ cells from neonatal mice resulted in higher frequency of p-STAT5^+^ T_FH_ cells than their adult counterparts.

Since IL-6 was expressed higher in immunized neonates along with heightened IL-2 signaling pathway involving T_FH_ cells, we investigated the link between IL-6 and IL-2 activity on neonatal T_FH_ cells. To assess the regulation of IL-2 activity when IL-6 is absent and when IL-6 is in excess, we immunized IL-6 KO mice in addition to repeating the IL-6 co-injection study [17]. The results of these two complementary experiments suggested an association between IL-6 and increased IL-2 production as well as elevated IL-2 receptor expression on T_FH_ cells. In neonatal mice, IL-2-expressing T_FH_ cell frequency was substantially increased when IL-6 was co-injected and decreased when IL-6 was absent (IL-6 KO mice). This inverse relationship was also present for the expression of IL-2 receptors and STAT5 phosphorylation as well as in T_FH_ generation. Corroborating the IL-6 co-injection and IL-6 KO mice immunization studies, CpG improved vaccine response where the diminished IL-6 and IL-2 production, together with reduced IL-2Rα and IL-2Rβ expression accompanied enhanced T_FH_, GC B cell and antibody responses.

At this point, it is not clear how IL-6 elicits a divergent function in regulating IL-2 production and IL-2 receptor expression on T_FH_ cells of adult and neonatal mice. Nevertheless, the unveiling of IL-6-mediated suppression of neonatal vaccine responses that involve enhanced IL-2 activity on T_FH_ cells has implications for the development of vaccines targeting early age. Adjuvants that improve vaccine responses in adults through enhanced production of IL-6 may not be suitable for neonates and infants. For example, SARS-CoV-2 mRNA vaccines contain lipid nanoparticles (LNP), which are shown to improve host immune response through the production of IL-6 and the expansion of T_FH_ response in adult mice [33; 39]. To our knowledge, efficacy of these vaccines in newborns and infants younger than 6 months of age have not been reported. If the mRNA-LNP vaccines stimulate IL-6 production in neonates also, these vaccines may not elicit protective antibodies in this age groups due to enhanced sensitization of T_FH_ cells to IL-2. Taken together, our findings support the tailoring of vaccines intended for early age based on the unique properties of this age group.

## Materials and Methods

### Mice

Wild-type and IL-6 KO (B6.129S2-Il6(tm1Kopf/J) mice with a C57BL/6 genetic background were purchased from Jackson Laboratory (Bar Harbor, Maine), bred, and kept in pathogen-free animal facilities in accordance with FDA Center for Veterinary Medicine guidelines. Neonatal (5- to 7-day-old) and adult (8- to 10-week-old) mice were used for immunization experiments. All animal procedures were approved by FDA’s Institutional Animal Care and Use Committee (Protocol 2017-48).

### Immunization

Tetanus toxoid conjugated type 14 pneumococcal polysaccharide (PPS14-TT) vaccine was manufactured as described [40]. PPS14-TT vaccine was emulsified with aluminum hydroxide [Al(OH)_3_] (Thermo Fischer, Waltham, MA). Aluminum hydroxide constituted 1/4^th^ of injection volume. For IL-6 co-injection experiments, PPS14-TT together with recombinant IL-6 (500 ng/adult and 100 ng/neonate (R&D systems, Minneapolis, MN) was emulsified with aluminum hydroxide. The adjuvant CpG 1826 (TCCATGA*CG*TTCCTGACGTT) was synthesized at FDA core facility. The CpG (10 μg per neonate mouse) containing PPS14-TT vaccine was emulsified with aluminum hydroxide by stirring for 30 minutes prior to injection. One and 0.5 μg of vaccines were injected in 150 μl and 30 μl volumes per adult and neonatal mice, respectively.

### Antibody for FACS analysis

Single-cell suspensions were prepared from splenocytes. Dead cells were stained by incubating cell suspensions with Zombie Aqua (BioLegend, San Diego, CA) diluted at 1:1000 dilution in PBS for 15 minutes at room temperature. Cells were washed and stained using FACS buffer containing 2% FBS, 0.5M EDTA in PBS. The following antibodies were used for surface staining at room temperature for 30 minutes: α-CD4 (BD Biosciences, Franklin Lakes, NJ, 1:100, GK1.55), α-B220 (BioLegend, 1:100, RA3-6B2), α-PD-1 (BD Biosciences, 1:100, J43 or BioLegend, 29F.1A12), α-CXCR5 biotin (BD Biosciences, 2G8), α-GL7 (BD Biosciences, GL-7), α-CD95 (BD Biosciences, J02), α-CD25 (BioLegend, 1:100, PC61), α-IL-2Rβ (BioLegend, 1:100, TM-β1), α-CD19 (BioLegend, 1:100, 6D5), α-CD3 (BioLegend, 1:100, 17A2), α-CD11c (BioLegend, 1:100, N418). To detect biotinylated CXCR5, cells were further incubated with streptavidin-BV-421 (BD Bioscience, 1:100) for 30 minutes at room temperature. For intracellular staining, samples were fixed with the FoxP3 Fix/Perm buffer set, following the manufacturer’s (ThermoFisher Scientific, eBioscience, Waltham, MA) instructions. Samples were then intracellularly stained with α-FoxP3 (BD Biosciences, 1:100, MF23 or BioLegend, 1:100, 150D), α-IL-2 (BD Biosciences, 1:100, JES6-5H4) or α-IL-6 (BioLegend, 1:100, MP5-20F3) antibodies for 30 minutes at room temperature. Flow cytometry data were acquired on Fortessa or Fortessa X20 flow cytometers (BD Biosciences) and analyzed using the FlowJo software v10.8.1 (FlowJo, Ashland, OR).

### In vitro stimulation of cells

For the assessment of IL-2 production, single-cell suspensions of splenocytes were stimulated with or without PMA (25 ng/ml, Sigma-Aldrich) and ionomycin (500 ng/ml, Invitrogen) in the presence of Brefeldin A (1:1000, Invitrogen) at 37°C for 4 h. After incubation, dead cells were stained with Zombie Aqua for 15 minutes at room temperature followed by surface marker staining. The cells were then fixed and permeabilized with FoxP3 Fix/Perm buffer set (ThermoFisher) and incubated with antibodies for IL-2 (BD Biosciences, 1:100, JES6-5H4) for 30 minutes at room temperature and analyzed in flow cytometry.

### Phospho-STAT5 FACS Analysis

Single-cell suspensions were incubated in culture media supplemented with 10% FBS, alone or with mouse recombinant IL-2 (R&D Systems, 50 ng/ml) for 15 minutes at 37°C. After washing with FACS buffer containing 1% FBS with 1mM EDTA, cells were first fixed with BD Cytofix fixation buffer (BD Bioscience) for 10 minutes at 37°C, and then permeabilized with pre-chilled BD Phosphoflow buffer III (BD Bioscience) for 10 minutes at 4°C. Cell surface antibodies and phospho-STAT5 (BD Bioscience, pY694 [clone 47]) antibody were incubated together in FACS buffer for 30 minutes at room temperature. To detect biotinylated CXCR5, cells were further incubated with streptavidin-BV-421 (BD Bioscience, 1:100) for 30 minutes at room temperature. For detection of T_FR_ cells, cells were washed and FoxP3 antibody was added to the permeabilization buffer (eBioscience) for 30 minutes at room temperature.

### Measurement of antibody titers against PPS14

For antibody measurement, 96-well plates were coated with purified PPS14 (ATCC, Manassas, Virginia) at 10 μg/ml in PBS (pH of 7) for 2 h at room temperature and then blocked for 1 h at room temperature with 5 % neonatal calf serum (Millipore Sigma, St. Louis, MO) in PBS. Serum samples (1:20 dilution) were serially diluted and 100 μl of diluted samples were transferred on coated plates for overnight incubation. After washing, wells were incubated with horseradish peroxidase-conjugated goat anti-mouse IgG-Fc or IgA antibody (Bethyl Laboratories, Waltham, MA) for 3 h at room temperature. For detection, 100 μl of KPL SureBlue TMB microwell peroxidase substrate (Seracare, Gaithersburg, MD) is added to the wells and incubated for 15-30 minutes followed by addition of stop solution (KPL TMB BlueSTOP solution, Seracare). The absorbance is measured at 450 nm.

### Statistical Analysis

Unpaired student’s t-test and One-Way ANOVA was used for all comparisons; data represented as mean +/− SEM are shown. P values <0.05 were considered statistically significant. *P<0.05, **P<0.01, ***P<0.001, ***P<0.0001.

## Supplemental Figures

### Supplemental Figure Legends

**Supplemental Figure 1.**
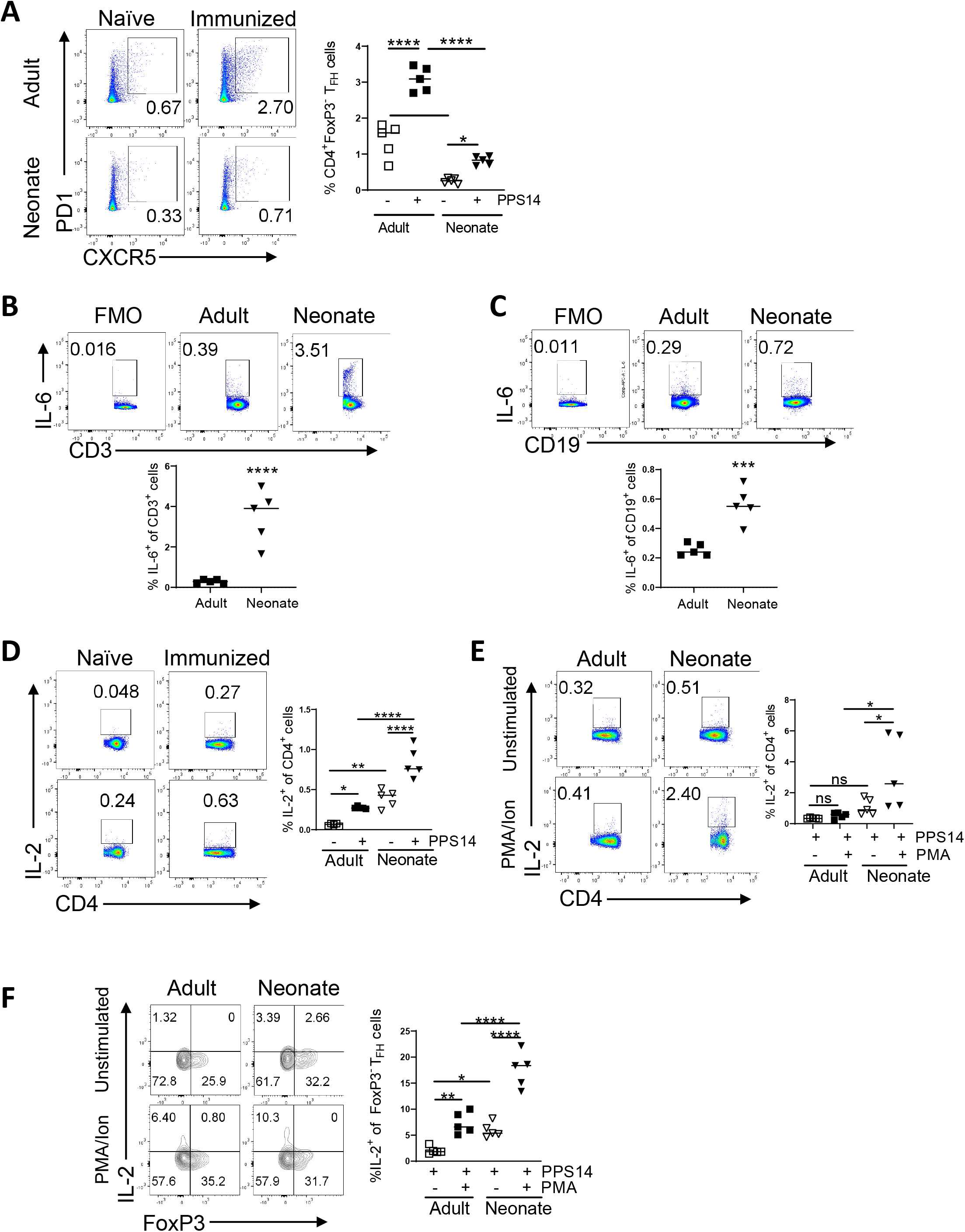
Adult (6- to 10-week-old) and neonatal (5- to 7-day-old) mice were immunized i.p. with PPS14-TT and splenocytes were analyzed by FACS. **(A)** Representative dot plots from immunized and unimmunized mice depict the percentages of T_FH_ (CXCR5^+^PD1^+^) cells pre-gated on CD4^+^FoxP3^−^ cells 7 dpi. Mean percentages of T_FH_ cells are plotted (n=5). **(B and C)**. Splenic cells were analyzed 24 hours post immunization. **(B)** Representative dot plots depict the percentages of IL-6^+^CD3^+^ cells. Mean percentages of IL-6^+^ cells among CD3^+^ cells are plotted (n=5). **(C)** Representative dot plots depict the percentages of IL-6^+^ cells pre-gated on CD19^+^ cells. Mean percentages of IL-6^+^ cells on CD19^+^ cells are plotted (n=5). Experiments were performed twice. **(D)** Representative dot plots depict the percentages of IL-2 expressing CD4^+^ cells from naïve and PPS14-TT immunized mice. Mean percentages of IL-2^+^ among CD4^+^ cells are plotted (n=5). **(E** and **F)** Splenocytes from immunized mice were in vitro stimulated with PMA/Ion for 4 h and cells were analyzed by FACS. **(F)** Representative dot plots depict the percentages of IL-2^+^ expressing CD4^+^ cells from unstimulated and PMA/Ion splenocytes. Mean percentages of IL-2^+^ among CD4^+^ cells are plotted (n=5). **(F)** Representative counter plots depict the percentages of IL-2-expressing Foxp3^+^ and Foxp3^−^ cells pre-gated on T_FH_ (CD4^+^CXCR5^+^PD1^+^) population. Mean percentages of IL-2^+^ cells among Foxp3^−^ T_FH_ cells are plotted (n=5). Experiments were performed twice.

**Supplemental Figure 2.**
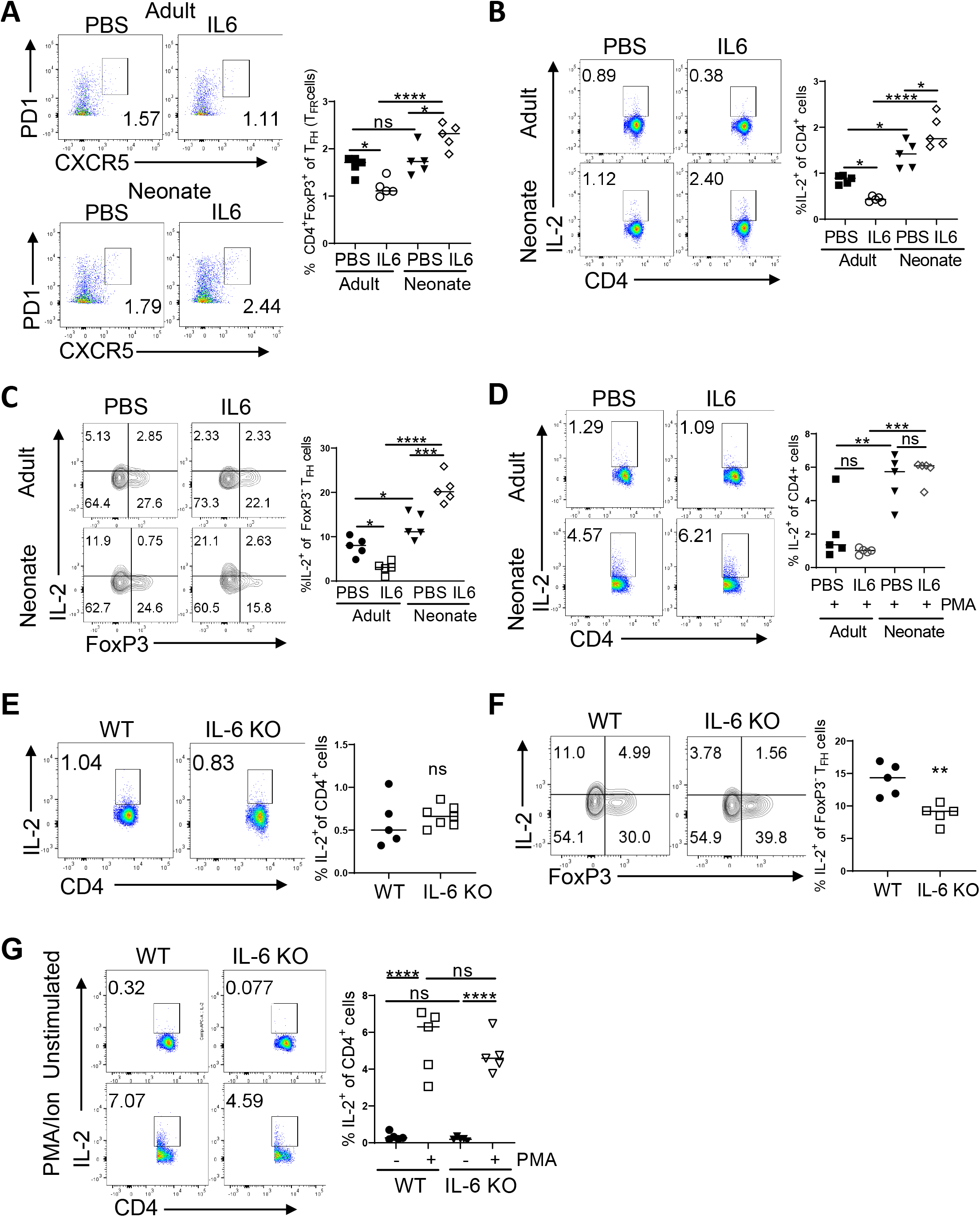
Adult (6- to 10-week-old) and neonatal (5- to 7-day-old) mice were immunized i.p. with PPS14-TT alone or PPS14-TT together with IL-6 and splenocytes were analyzed by FACS. **(A)** Representative dot plots show CXCR5^+^PD1^+^ T_FR_ cells pre-gated on CD4^+^FoxP3^−^ population in immunized mice. Mean percentages of T_FR_ cells are plotted (n=5). **(B)** Representative dot plots depict the percentages of IL-2 expressing CD4^+^ cells from immunized mice. Mean percentages of IL-2^+^ among CD4^+^ cells are plotted (n=5). **(C)** Representative counter plots depict the percentages of IL-2-expressing Foxp3^+^ and FoxP3^−^ cells pre-gated on T_FH_ (CD4^+^CXCR5^+^PD1^+^) population. Mean percentages of IL-2^+^ cells among FoxP3^−^ T_FH_ cells are plotted (n=5). **(D)** Splenocytes from immunized mice were in vitro stimulated with PMA/Ion for 4 h and cells were analyzed by FACS. Representative dot plots depict the percentages of IL-2^+^ expressing CD4^+^ cells from stimulated splenocytes (n=5). Experiments were performed three times. **(E – G)** Neonatal (5- to 7-day-old) wild-type (C57BL/6J) and IL-6 KO mice were immunized i.p. with PPS14-TT and splenocytes were analyzed by FACS. **(E)** Representative dot plots depict the percentages of IL-2 expressing CD4^+^ cells from immunized mice. Mean percentages of IL-2^+^ among CD4^+^ cells are plotted (n=5). **(F)** Representative counter plots depict the percentages of IL-2-expressing Foxp3^+^ and Foxp3^−^ cells pre-gated on T_FH_ (CD4^+^CXCR5^+^PD1^+^) population. Mean percentages of IL-2^+^ cells among Foxp3^−^ T_FH_ cells are plotted (n=5). **(G)** Representative dot plots depict the percentages of IL-2^+^ expressing CD4^+^ cells from unstimulated and PMA/Ion splenocytes. Mean percentages of IL-2^+^ among CD4^+^ cells are plotted (n=5). Experiments were performed three to four times.

**Supplemental Figure 3.**
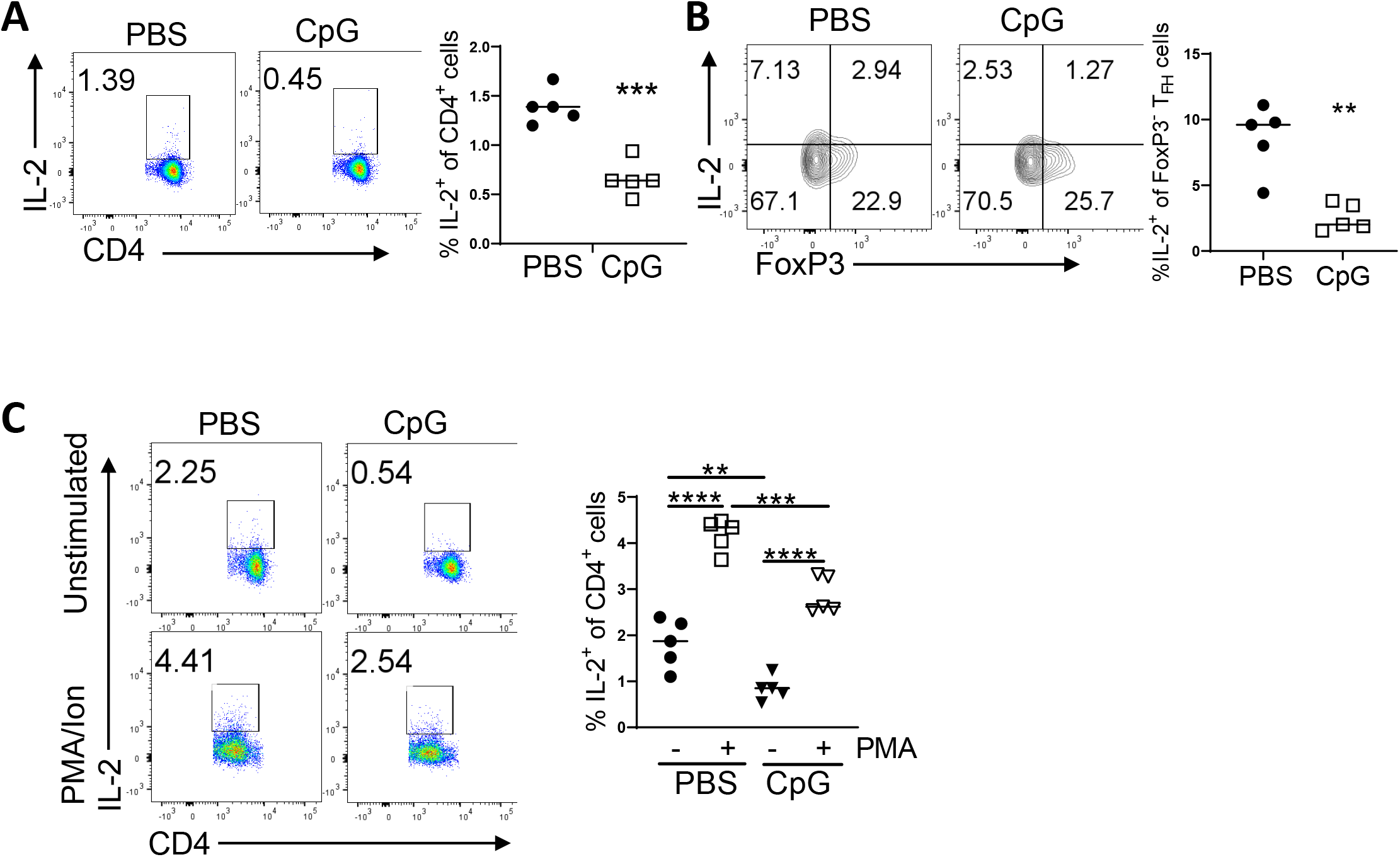
C57BL/6J (5- to 7-day-old) mice were immunized i.p. with PPS14-TT (PBS) or PPS14-TT + CpG (CpG) and splenocytes were analyzed by flow cytometry at 7 dpi. **(A)** Representative dot plots depict the percentages of IL-2 expressing CD4^+^ cells from immunized mice. Mean percentages of IL-2^+^ among CD4^+^ cells are plotted (n=5). **(B)** Representative counter plots depict the percentages of IL-2-expressing Foxp3^+^ and Foxp3^−^ cells pre-gated on T_FH_ (CD4^+^CXCR5^+^PD1^+^) population. Mean percentages of IL-2^+^ cells among Foxp3^−^ T_FH_ cells are plotted (n=5). **(D)** Representative dot plots depict the percentages of IL-2^+^ expressing CD4^+^ cells from unstimulated and PMA/Ion splenocytes. Mean percentages of IL-2^+^ among CD4^+^ cells are plotted (n=5). Experiments were performed twice.

